# Combating pancreatic cancer with ovarian cancer cells

**DOI:** 10.1101/2022.09.01.506158

**Authors:** Xiao Lin, Chunmei Cui, Qinghua Cui

**Affiliations:** Department of Biomedical Informatics, School of Basic Medical Sciences, Peking University, 38 Xueyuan Rd, Beijing, 100191, China

**Author notes:** To whom the correspondence should be addressed: Dr. Qinghua Cui, Ms. Xiao Lin, Dr. Chunmei Cui. Equal contribution (Xiao Lin, Chunmei Cui).

**Keywords:** Pancreatic cancer, cancer therapy, ovarian cancer

## Abstract

With overall five-year survival rate less than 10%, pancreatic cancer (PC) represents the most lethal one in all human cancers. Given that the incidence of PC is still increasing and current cancer treatment strategies are often inefficacious, its therapy is still a huge challenge. Here, we first revealed ovarian serous carcinoma is mostly anti-correlated with pancreatic cancer in gene expression signatures. Based on this observation, we proposed that ovarian cancer cells could defense PC. To confirm this strategy, we first showed that ovarian cancer cell SKOV3 can significantly inhibit the proliferation of pancreatic cancer cell SW1990 when they were co-cultured. We further validated this strategy by an animal model of pancreatic cancer xenografts. The result showed that the injection of SKOV3 significantly inhibits pancreatic cancer xenografts. Moreover, we found that SKOV3 with transgenic African elephant *TP53* gene further enhances the therapeutic effect. RNA-sequencing analysis revealed that the ovarian cancer cell treatment strikingly induced changes of genes being involved in pancreas function and phenotype (e.g. enhancing pancreas function, pancreas regeneration, and cell adhesion) but not immune and inflammation related functions, suggesting that the proposed strategy is different from immunotherapy and could be a novel strategy for cancer treatment.

## Introduction

Pancreatic cancer, an aggressive disease that typically diagnosed at an advanced stage, is the seventh leading cause of cancer death worldwide according to global cancer statistics in 2018 ^1, 2^. Moreover, with a persistently increasing incidence, pancreatic cancer would become the second leading cause of cancer-related mortality by 2030 ^3^.

At present, the main therapies for pancreatic cancer include surgery, chemotherapy, radiation therapy, interventional therapy and immunotherapy. Surgical resection with adjuvant systemic chemotherapy currently provides the only chance for improving survival of patients with pancreatic cancer ^4^. However, more than 80% of pancreatic cancer patients are in the middle and late stages when they are diagnosed and thus lose the opportunity for surgery. Chemotherapy is the main treatment for patients with unresectable advanced pancreatic cancer. The clinical routine chemotherapy for pancreatic cancer includes gemcitabine monotherapy, gemcitabine combined with nab-paclitaxel and FOLFIRINOX regimen (fluorouracil, leucovorin, irinotecan, and oxaliplatin) ^5, 6, 7^. Among them, liposomal irinotecan is suitable for combination with fluorouracil and folinic acid in patients with advanced pancreatic cancer refractory to gemcitabine chemotherapy (second-line therapy). The combination chemotherapy program developed in the past ten years can help patients with advanced and metastatic pancreatic cancer have short-term partial remission or stable disease, but almost all patients will eventually relapse ^8, 9^. Recent research on the genome of pancreatic cancer has promoted the development of targeted therapy ^10^. But it is only suitable for a small number of pancreatic cancer patients and only improves limited survival time. For immunotherapies, it is known that ipilimumab (cytotoxic T-cell antigen-4 inhibitor) has a good curative effect on advanced melanoma ^11^ and PD-L1 monoclonal antibody (immune checkpoint inhibitor) is sensitive to melanoma ^12^, non-small cell lung cancer ^13^, Hodgkin’s lymphoma ^14^ and urothelial carcinoma ^15^. However, limited studies have confirmed these immunotherapies for advanced pancreatic cancer ^16, 17^. CAR-engineered T cells (CAR-T cells) therapeutics have shown promising outcomes in treating haematological malignancies ^14^. However, clinical trials showed its efficiency is limited for patients suffering from solid tumors ^18–20^. Given the difficulty of diagnosing pre-pancreatic cancer and the shortcomings of the current treatment strategies, the 5-year overall survival rate of pancreatic cancer patients is less than 10% ^21^ and the pancreatic cancer represents the most lethal one in all human cancers. Therefore, new strategies of treatment are urgently needed to improve the survival of pancreatic cancer patients.

Reverse-transcriptomics represent one novel and popular technique to explore novel agents for disease treatment through identifying possible agents whose gene expression signature is significantly anti-correlated with that of the candidate disease ^22^. In this study, we comprehensively screened possible agents whose gene expression signatures are anti-correlated with pancreatic ductal adenocarcinoma cancer (PDAC), the most common pancreatic cancer using DrugMine, a reverse-transcriptomics based computational tool. As a result, the human ovarian serous carcinoma is one of the top candidate agents, suggesting that ovarian cancer cell transplantation could be used for pancreatic cancer therapy. We next confirmed this strategy by cell experiment and series of animal experiments. Finally, analysis of RNA-sequencing data revealed ovarian cancer cell induced gene signature is mainly involved in pancreas function and phenotype in pancreatic tumors, such as enhancing pancreas function, pancreas regeneration, and cell adhesion. The immune and inflammation related functions and pathways are not significant. The above results suggest that the proposed strategy is different from immunotherapy, which then further suggest that the proposed method would represent a novel and general strategy for cancer therapy, that is, combating cancer with cancer (CCC) therapeutics.

## Methods & Materials

### Animals

BALB/c nude male mice (6-8 week-old, weighing 16-20g) of SPF-class were purchased from Huafukang Animal Experiment Co., Ltd (Beijing, China). The mice were housed under in a temperature-controlled environment with a 12/12 h light/dark cycle with standard diet and water *ad libitum*. All animal procedures were handled according to the guidelines of the laboratory animal care (NIH publication no.85Y23, revised 1996) and approved by the Animal Care and Use Committee of the Peking University Health Science Center.

### Cell Lines and Lentivirus

Human pancreatic cancer cell line SW1990 was purchased from ATCC Co., Ltd (USA) and human ovarian cancer cell line SKOV3 was provided by Prof. Lixiang Xue of Peking University Third Hospital. Cell lines were maintained in RPMI 1640 (Gbico, USA) supplemented with 10% fetal bovine serum (Gibco, USA). Cells were cultured at 37°C in a humidified incubator containing 5% CO_2_.

Lentivirus vectors containing the GFP tag, GFP tag and *TP53* gene of *Loxodonta africana* were constructed, respectively. Lentivirus production was completed by the Genechem Company (Shanghai, China). SKOV3 cells were infected with the concentrated virus at a multiplicity of infection of 5 in the presence of HitransG A for 12 h. After 12 hours, the supernatant was replaced with fresh medium. GFP-positive SKOV3 cells were sorted by flow cytometry, and expression of *TP53* in the infected cells evaluated by q-PCR.

### Cell experiment

10^5^ SKOV3^GFP^ cell and 10^5^ SW1990 cell / per well were co-cultured in 6-well plate, in which 2×10^5^ SW1990 cells / per well were planted in 6-well plate as a control. After 48 hours, 3 fields were selected from each well to count the total cells in the bright field and GFP cells (SKOV3 cells) in the fluorescence field, then the number of SW1990 cells was equal to the number of total cells minus the number of GFP cells (SKOV3 cells). The relative cell proliferation rate of SW1990 was calculated with a formula: (the cell number of SW1990 × 2 in co-cultured group) / (the cell number of SW1990 in control group) × 100%.

### Quantitative real-time PCR assay

Total RNA from SKOV3 cells was isolated by using TriQuick reagent (R1100, Solarbio) and 5 μg RNA was reverse-transcribed for preparing cDNA by a commercial RT kit (Transgene, AH341). Real-time PCR was performed using the AriaMx Real-Time PCR System with Top Green PCR Master Mix (AQ131-01, TransGen Biotech) and the following primer: *TP53* (F:TTTCACCCTTCAGATCCGTG; R:GACTGTCCCTTCTTAGACTTCG), *GAPDH*(F:CTC CTCCACCTTTGACGCTG, R: TCCTCTTGTGCTCTTGCTGG), *GAPDH* was used as the internal reference.

### Establishment of subcutaneous transplantation tumor model

To produce SW1990 donor tumors, SW1990 cell suspension of logarithmic growth phase in the total volume of 0.2 mL was inoculated subcutaneously into the right posterior axillary line of nude mice. When the tumor grows to a certain extent, the tumor was aseptically peeled off and cut into small tissue pieces of 2×2×2 (mm^3^), and the tumor tissue pieces were inoculated into the right axilla of nude mice subcutaneously through a trocar to complete the subcutaneous pancreatic cancer model. The tumor volume was calculated with a standard formula: width^2^ × length × 0.5. At the end of the experiment, the xenografts were weighted, excised, fixed in formalin or frozen at −80°C for further experiments.

### Screening possible agents to defense pancreatic cancer

To explore possible agents for pancreatic cancer defense, we first got the gene-level expression profiles of 28 PDAC specimens and 2 normal pancreatic tissues from donors (GEO accession number: GSE56560) ^23^. Genes with fold change (FC) value ≥1.50 or ≤0.67 are taken as the gene signature of PDAC. And then the reverse-transcriptomics based computational tool DrugMine from Co., Ltd of Beijing JeaMoon Technology was used to explore possible agents (including small molecules, biological molecules, and traditional Chinese medicine etc) to defense pancreatic cancer. The hypothesis of DrugMine is that if the gene signature of some agent is anti-correlated with that of PDAC, then this agent will be predicted to be able to defense pancreatic cancer. In addition, Spearman’s correlation was also performed in DrugMine based on the whole FC profiles.

### RNA-sequencing

RNA sequencing was performed using RNA extracted from tumors in normal saline control (NSCT) and SKOV3 high dose (HD) group by using TriQuick reagent (R1100, Solarbio). Each group consists of 3 biological replicates. RNA degradation and contamination was monitored on 1% agarose gels. RNA purity was checked using the NanoPhotometer^®^ spectrophotometer (IMPLEN, CA, USA). RNA integrity was assessed using the RNA Nano 6000 Assay Kit of the Bioanalyzer 2100 system (Agilent Technologies, CA, USA). RNA-sequencing was performed according to the standard protocol of Novogene Corporation Inc. The RNA-sequencing data is publicly available at GEO (GSE212173). Differential expression analysis was performed using the DESeq2 R package (1.16.1).

### Functional and pathway analysis

To explore possible mechanisms of the CCC therapeutics for combating pancreatic cancer with ovarian cancer cells, we applied DAVID Bioinformatics ^24^ and the weighted functional and pathway analysis tool WEAT ^25^, through which here we analyzed a number of functional/pathway categories, including ARCHS4 Kinases Coexp, ARCHS4 TFs Coexp, BioPlanet, BioPlex, ClinVar 2019, DisGeNET, DSigDB, GO biological process, GO cellular component, GO molecular function, GWAS Catalog 2019, Human Gene Atlas, Human Phenotype Ontology, Jensen COMPARTMENTS, Jensen DISEASES, KEGG, MSigDB Hallmark 2020, Pfam Domains 2019, Rare Diseases AutoRIF Gene Lists, RNA Seq Disease Gene and Drug Signatures from GEO, and TRANSFAC and JASPAR PWMs. Genes with fold change (FC) >=2.0 or <=0.5 between the pancreatic cancer xenografts with and without SKOV3 treatment as the differentially expressed genes (DEGs).

### Quantification and Statistical Analysis

All data was expressed as the mean ± SD. R (version 4.2) was used for all statistical analyses. For a comparison between two groups, the T-test was used. For comparisons between three or more groups, the one-way ANOVA (one-tailed) was used.

## Result

### Top agents predicted to reverse gene signature of pancreatic cancer

We screened the candidate agents whose gene signatures are anti-correlated with that of pancreatic cancer using DrugMine. As a result, Figure 1A shows the top ten agents with most significant odds ratio (OR) value by Fisher’s exact test and most significant Spearman’s correlation coefficient (Rho). The p-values of both tests for all agents are almost zero, indicating that their induced gene expression changes are significantly anti-correlated with that of pancreatic cancer, and further suggests that these agents could defense pancreatic cancer. Moreover, strikingly, we noted that ovarian serous carcinoma is among the top ten agents, indicating that it is anti-correlated with pancreatic cancer in gene expression changes. It is well known that cancers mostly share common pathways and mechanisms during their formation and development. However, here clear discrepancy was revealed between ovarian serous carcinoma and pancreatic cancer. This finding hinted that the opposite mechanisms could exist between the two cancers and thus suggested a possible novel therapeutics, that is, killing pancreatic cancer with ovarian cancer cells. Based on the above observation, we proposed a novel therapeutics for pancreatic cancer treatment as shown in Figure 1B, that is, injection of ovarian cancer cells into pancreatic tumor could inhibit pancreatic cancer.

**Figure 1.**
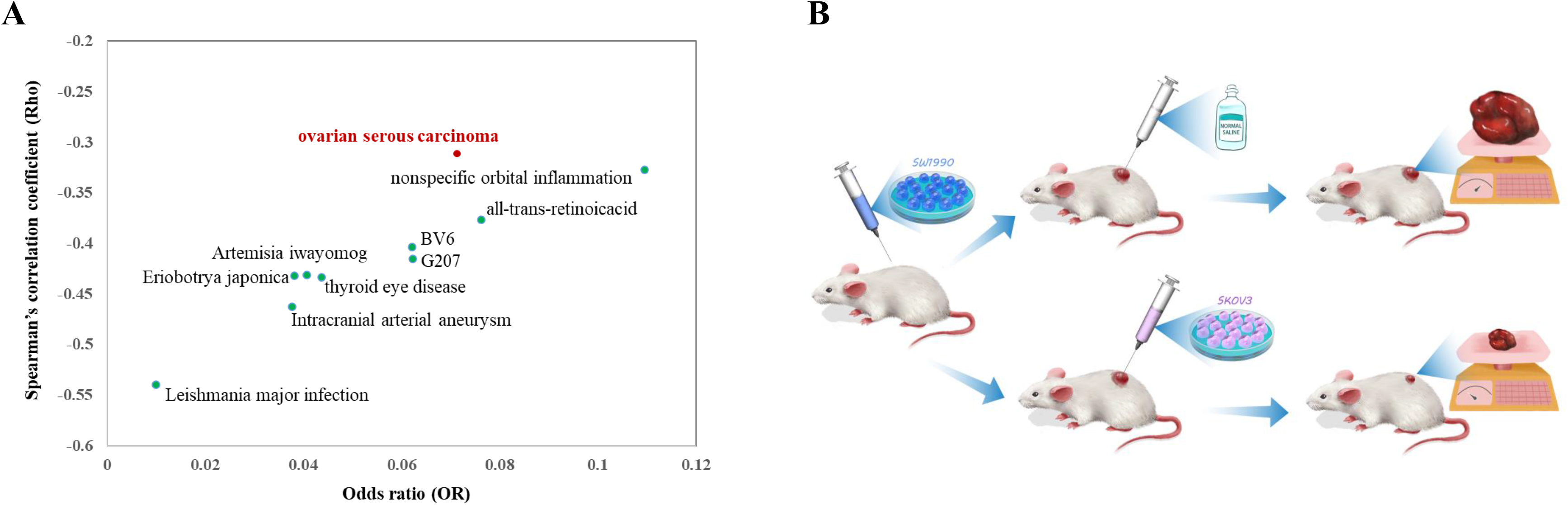
The top ten agents with least odds ratio (OR) value by Fisher’s exact test and least Spearman’s correlation coefficient (Rho) with that of pancreatic cancer (**A**) and the work flow of the proposed therapeutics of combating pancreatic cancer with ovarian cancer cell (**B**).

### Injection of ovarian cancer cell SKOV3 inhibited pancreatic tumor development

Firstly, we tested whether ovarian cancer cell SKOV3 could inhibit pancreatic cancer cell SW1990 *in vitro*, so we sorted and purified SKOV3 cells which were transfected with lentivirus carrying the GFP tag through GFP fluorescence by flow sorting. By co-culturing the SKOV3 cells with SW1990 cells, it was found that SKOV3 could reduce the proliferation rate of SW1990 by about 37% and showed a significant inhibition (Figure 2A, p-value = 0.0036, T-test), suggesting that ovarian cancer cell indeed can inhibit pancreatic cancer cell *in vitro*. In order to confirm whether SKOV3 cells can inhibit pancreatic cancer *in vivo*, we first established subcutaneous pancreatic cancer xenografts by subcutaneous injection of SW1990 cells, then followed by intervention with low dose (LD, 3×10^5^ cells) or high dose (HD, 6×10^5^ cells) SKOV3 cells twice a week through intratumoral injection. Equivalent normal saline solution (NSCT) was given twice a week by intratumoral injection as blank control. Gemcitabine was used as a positive control in the intervention group (100mg/kg, twice a week by intraperitoneal injection). The mice were euthanized and the tumors were harvested after 11 days of intervention (Figure 2B). When compared with the NSCT group, we found a significant decrease of tumor weight in Gemcitabine group and the SKOV3 HD group after 11 days of treatment (Figure 2C, Gemcitabine vs. NSCT, p-value = 0.0015; SKOV3 HD vs. NSCT, p-value = 0.0278, one-way ANOVA test). When compared with the NSCT group, we found a dramatically decreased tumor volume in the Gemcitabine group, SKOV3 LD group and the SKOV3 HD group after 11 days of treatment (Figure 2D, Gemcitabine vs. NSCT, p-value = 0.0022; SKOV3 HD vs. NSCT, p-value = 0.0098; SKOV3 LD vs. NSCT, p-value = 0.0211, one-way ANOVA test). Not only that, the tumor volume was detected on the 4th and 7th day after treatment, which was significantly reduced compared with the NSCT group (Supplementary File S1). In addition, there was no significant difference in body weight change among the four groups (Figure 2E), but there was a tendency in slowing body weight loss in the SKOV3 HD group. The above results suggest that the ovarian cancer cell indeed can inhibit the growth of pancreatic tumors *in vivo*.

**Figure 2.**
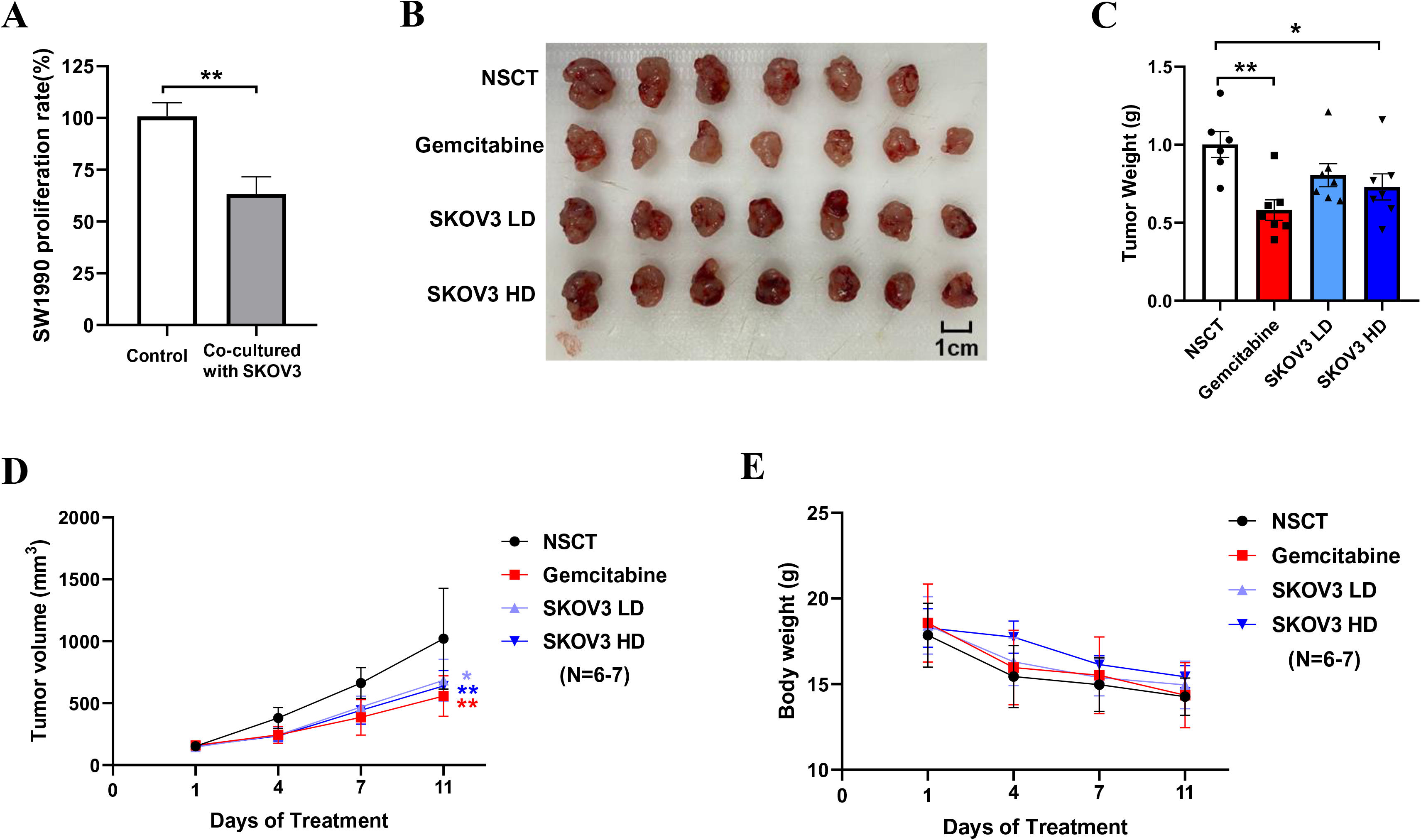
The results of the cell experiment and animal experiment based on SKOV3^GFP^ cells and wild-type SKOV3 cells. The inhibition of SKOV3 cell on SW1990 cell *in vitro*(**A**). After 11 days of treatment, the mice were euthanized for analysis of tumors. The images of harvested tumors (**B**). Tumor weights are shown as means ± SD (**C**). *n* =6-7, ^*^*p* < 0.05, ^**^*p* < 0.01. Tumor growth curve during intervention (**D**). The points and bars represent means ± SD. *n* =6-7, ^*^*p* < 0.05, ^**^*p* < 0.01. Body weight changes of mice during intervention (**E**).

### Potential mechanisms of the proposed CCC therapeutics

As we reported above, the proposed CCC therapeutics indeed inhibited the growth of pancreatic cancer through injection of ovarian cancer cells. As we described above, this therapeutics was proposed based on the observation of gene expression change of ovarian cancer is anti-correlated with that of pancreatic cancer (Rho=−0.31, p-value= 0, Figure 3A). To confirm whether injection of ovarian cancer cell reverse the gene expression change of pancreatic cancer, we identified the gene expression profiles of subcutaneous xenotransplanted tumors with (the high dose group) or without injection of ovarian cancer cells by RNA-sequencing. As expected, the ovarian cancer cell induced gene expression change (with vs. without ovarian cancer cell treatment) is indeed anti-correlated with that (pancreatic cancer vs. normal pancreas) of pancreatic cancer (Rho=−0.19, p-value= 2.07e-95, Figure 3B). This result means that the ovarian cancer cell treatment indeed statistically reverses the gene expression signature of pancreatic cancer and thus together with the cell and animal results confirmed the efficacy of the CCC therapeutics for pancreatic cancer. Given that the proposed CCC therapeutics is based on injection of cancer cells, however, people would regard it as one kind of immunotherapy. To confirm whether it belongs to immunotherapy or not and to explore whether it represents some novel therapeutics, we next performed function and pathway enrichment analysis for the DEGs of pancreatic tumors with and without ovarian cancer cell treatments using tools of David Bioinformatics and WEAT. Firstly, we investigated the enriched subcellular locations of the DEGs. As a result, both DAVID Bioinformatics and WEAT analysis revealed that the DEGs are mostly enriched in the extracellular space/region and cell membrane/surface (Figure 3C), which is consistent with the enriched functions of signaling, ion transport, extracellular matrix organization, and pancreatic secretion (Figure 3D).

**Figure 3.**
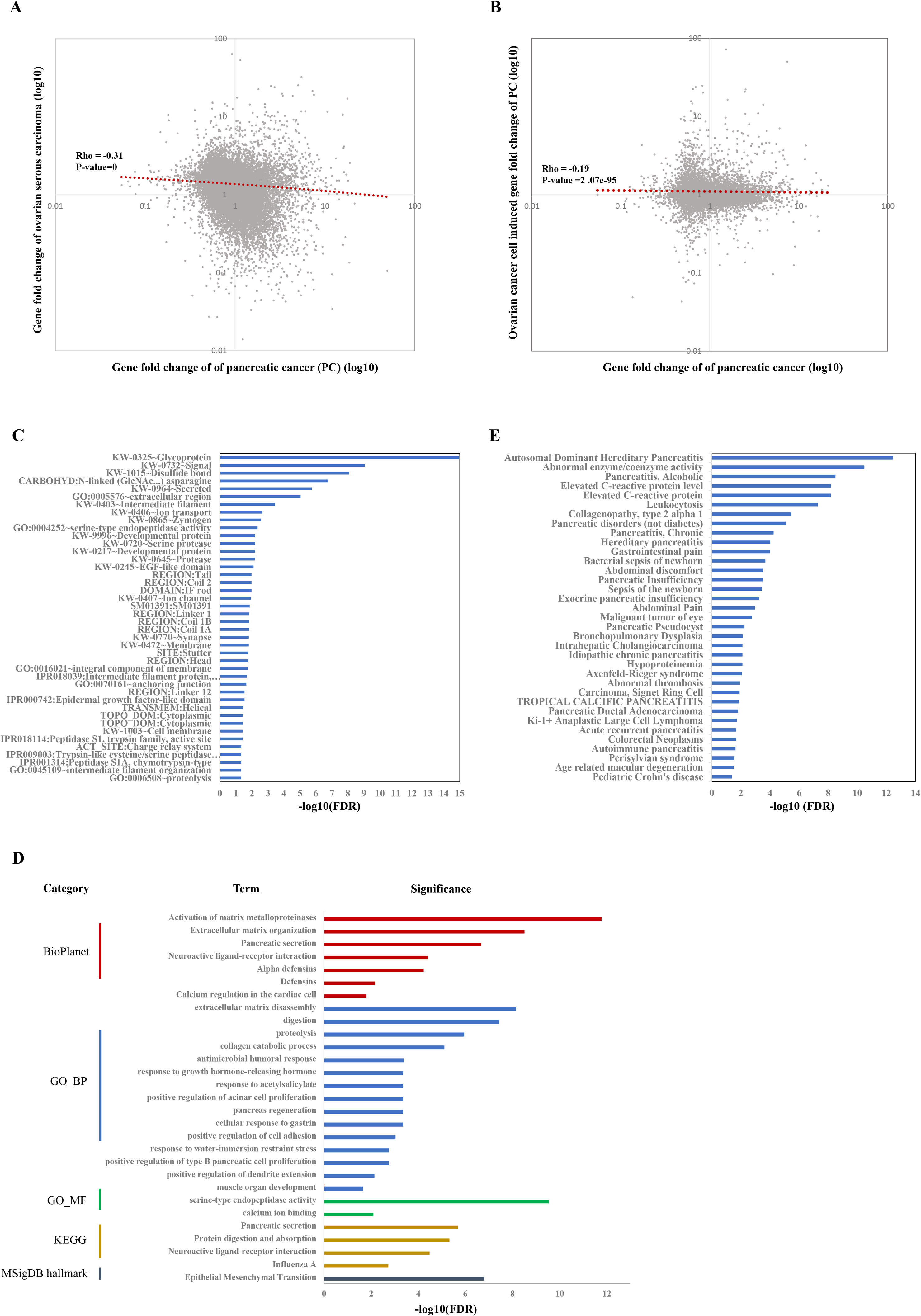
Function, pathways, and targets induced by SKOV3 cell treatment. Spearman’s correlation between gene fold change of ovarian serous carcinoma and gene fold change of pancreatic cancer (**A**). Spearman’s correlation between gene fold change of pancreatic cancer induced by ovarian cancer cell SKOV3 and gene fold change of pancreatic cancer (**B**). The enrichment result of xenograft with or without injection of SKOV3 cells by using David (**C**). The enrichment result of xenograft with or without injection of SKOV3 cells by using WEAT (**D**). The enriched diseases in DisGeNET of xenograft with or without injection of SKOV3 cells (**E**).

Next, investigating the phenotype and disease of these DEGs involved in may provide critical insights for exploring the mechanisms of the CCC therapeutics. As a result, pancreatic-related diseases are among the top terms (Supplementary File S2). For example, the term of pancreatitis especially hereditary/pediatric pancreatitis is one of the top significant terms in all categories (ClinVar, DisGeNET, GWAS, Human Phenotype Ontology, Jensen DISEASES, and rare disease). In addition, the DEGs are also enriched in a number of other pancreatic-related disease, such as alcoholic pancreatitis, non-diabetic pancreatic disorders, chronic pancreatitis, pancreatic insufficiency, exocrine pancreatic insufficiency, pancreatic pseudocyst, idiopathic chronic pancreatitis, pancreatic ductal adenocarcinoma, acute recurrent pancreatitis, and autoimmune pancreatitis in the DisGeNET category (Figure 3E), and pancreatic steatorrhea in the Jensen DISEASES category. The above results suggest that the ovarian cancer cell treatment strikingly induced changes of genes being involved in pancreas function and phenotype in pancreatic tumors.

Given that the pancreas-related disease including pancreatitis is the top term related with the DEGs, it will be interesting whether the DEGs contribute to these phenotypes through inflammation or immune related functions and pathways. For doing so, we performed functions and pathways analysis (Figure 3D). Strikingly, biological processes such as protein digestion and absorption, proteolysis, response to growth hormone-releasing hormone, pancreas regeneration, positive regulation of cell adhesion, and positive regulation of type B pancreatic cell proliferation were mostly significant, suggesting that the CCC therapeutics could work by enhancing pancreas function, pancreas regeneration and cell adhesion through secreting signaling molecules to trigger proteolysis but not through inflammation and immune related functions and pathways. Moreover, molecular function (MF) analysis showed that the proteolysis could be mediated by serine-type endopeptidase but not other peptidase. Interestingly, the only significant term in MSigDB hallmark category is ‘Epithelial Mesenchymal Transition’ (EMT, FDR=1.55e-7), suggesting that another possible mechanism of the CCC therapeutics is that the ovarian cancer cell could block the metastasis of pancreatic cancer by inhibiting its EMT process. Specifically, immune-related functional or pathway terms are not identified as significance, suggests that the CCC therapeutics could not belong to immunotherapy.

We next try to explore genes or targets contributing to or being highly related to these DEGs, including categories of kinase, transcription factor (TF), BioPlex Interactome, and Pfam domains. As a result, 81 kinases and 109 TFs were identified to be mostly co-expressed with the DEGs (Supplementary File S3 & S4). It should be noted that NR5A2 (nuclear receptor subfamily 5 group A member 2) is the top 1 significant TF. Interestingly, it was reported that NR5A2 has critical roles in restraining inflammation in normal mouse pancreas and is considered to be highly relevant to human pancreatitis and pancreatic cancer ^26^. In BioPlex analysis, there 7 significant genes highly connected with these DEGs, including GDF11, GSTT1, FN1, ITGA4, TUBB1, TUBB3, and HSPA5 (Supplementary File S5). All of the 7 genes have critical roles in carcinogenesis, for example, it is known that FN1 (fibronectin 1) is involved in cell adhesion and migration processes including metastasis, and thus is associated with pancreatic cancer prognosis and survival ^27^. For the Pfam Domain analysis, the only significant term is ‘Trypsin’ (FDR=4.72e-8), an enzyme produced in pancreas.

### African elephant *TP53* transgenic SKOV3 cells enhanced the inhibitory effects of pancreatic tumor

In the above section, we confirmed the efficacy of the CCC therapeutics on pancreatic cancer through injection of the SKOV3 ovarian cancer cells. It is then interesting to investigate whether gene engineering the ovarian cancer cell can further enhance the efficacy or not. It is known that *TP53* (tumor protein 53) is a crucial tumor suppressor gene, mutated in majority of human cancers^28^. Interestingly, the multiple copies of *TP53* and the enhanced p53-mediated cell apoptosis observed in African elephants may have evolved to offer such cancer protection^29^. Therefore, we transferred *TP53* gene of African elephant into SKOV3 cell to investigate if it can enhance the inhibition of the growth of pancreatic cancer.

We sorted and purified SKOV3 cells which were transfected with lentivirus containing GFP tag and *TP53* gene through GFP fluorescence by flow sorting, and detected the expression of *TP53* by q-PCR. The results of q-PCR also showed that the expression of *TP53* gene was significantly increased after transfection in SKOV3 cells (Figure 4A).

**Figure 4.**
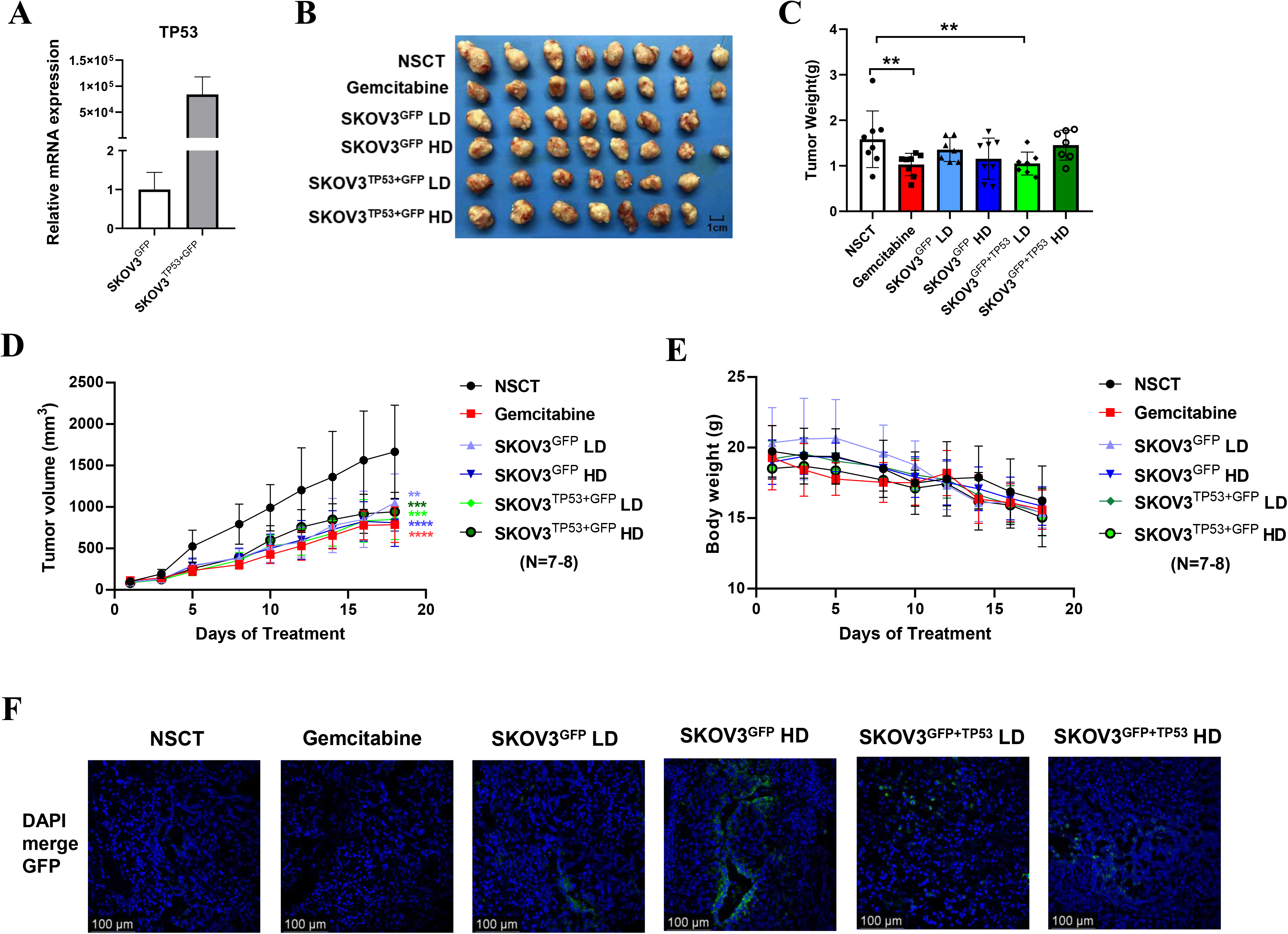
The results of the animal experiment based on African elephant *TP53* transgenic SKOV3 cells. The q-PCR analysis showed the mRNA expression levels of *TP53* in SKOV3 cell after the transfection of lentivirus (**A**). After 18 days of treatment, the mice were euthanized for analysis of tumors. The images of harvested tumors (**B**). Tumor weights are shown as means ± SD (**C**). n =7-8, **p < 0.01. Tumor growth curve during intervention (**D**). The points and bars represent means ± SD. n =7-8, **p < 0.01, ***p < 0.001, ****p < 0.0001. Body weight changes of mice during intervention (**E**). Representative images of xenograft of each group in fluorescence confocal (**F**). Nuclei were stained with DAPI (blue). Bars =100 μm.

Similar to the work flow of the previous animal experiments, we upped the dose and frequency of cell injection, and followed by intervention with low dose (LD, 6×10^5^ cells) or high dose (HD, 1.2×10^6^ cells) SKOV3^GFP^ or SKOV3^TP53+GFP^ cells every two days through intratumoral injection. As shown in Figure 4B, the mice were euthanized and the tumors were harvested after 18 days of intervention. As a result, the tumor weight in the Gemcitabine group and the SKOV3^TP53+GFP^ LD had a significant decrease (Figure 4C, Gemcitabine group vs. NSCT, p-value = 0.016; SKOV3^TP53+GFP^ LD vs. NSCT, p-value = 0.025, one-way ANOVA test). Comparing with the NSCT group, xenograft tumors grown in both the SKOV3^GFP^ group and the SKOV3^TP53+GFP^ group of low dose or high dose had a smaller mean size (Figure 4C). The tumor volume was significantly smaller and significantly different from the NSCT group at each measurement after intervention. Furthermore, there was no significant difference in body weight change among the four groups (Figure 4E), but, early in the intervention, treatment with SKOV3^GFP^ or SKOV3^TP53+GFP^ cells can slow down the weight loss. In addition, as SKOV3 is also a rapidly proliferating cancer cell, we were concerned that the injected SKOV3 cell might develop into ovarian cancer in the pancreatic tumor. Therefore, we observed the green fluorescence under confocal microscope after frozen section of xenograft tumor. The result showed that no ovarian cancer was formed in the pancreatic tumor after SKOV3^GFP^ and SKOV3^TP53+GFP^ injection, but only some scattered GFP fluorescence signals were detected (Figure 4F). The above results indicate that the CCC therapeutics combined with genetic engineering may further enhance the treatment effect.

## Discussion

Pancreatic cancer is a common malignant tumor of the digestive system. Due to its low surgical resection rate and poor sensitivity to various therapeutics such as chemotherapy and radiotherapy, it leads to a very low 5-year survival rate and represents the most lethal cancer. Therefore, bringing novel therapeutics to patients with pancreatic cancer has been a long-standing and challenging problem. In this study, we first predicted the candidate agents whose gene signatures are anti-correlated with that of pancreatic cancer by reverse-transcriptomics. As a result, we found ovarian cancer is among the top candidate agents, which hints that ovarian cancer cells may be used to defense pancreatic cancer. We next confirmed this hypothesis using cell and animal experiments. As a result, the ovarian cancer cell SKOV3 can indeed significantly inhibit the proliferation of the pancreatic cancer cell SW1990 and the growth of pancreatic tumors. Moreover, the RNA-seq analysis of subcutaneous xenotransplanted tumors with or without injection of SKOV3 cells showed that the gene expression change induced by ovarian cancer cell treatment is indeed anti-correlated with that of pancreatic cancer. Moreover, we showed that the transgenic *TP53* gene of African elephant in SKOV3 cell can further enhance the treatment effects. Together, our results showed the feasibility of the CCC or genetic engineering CCC therapeutics.

It is widely known that in epithelial cancers, including pancreatic cancer, EMT is associated with the three major steps of cancer development: invasion, dissemination and metastasis ^30, 31^. EMT change is triggered in a number of distinct molecular processes including the expression of specific cell-surface proteins and the activation of transcription factors (TFs). EMT-TFs includes SNAI and SNAI2 superfamily, two ZEB factors (ZEB1 and ZEB2) ^30^ and other TFs, like Prrx1, Sox4 and Sox9, Klf4 and members of the AP-1 (Jun/Fos) family ^32^. In our results, function and pathway analysis of DEGs in xenografts with or without injection of SKOV3 cells revealed several pathways that are important in the development of cancer, including epithelial mesenchymal transition (EMT), activation of matrix metalloproteinases, extracellular matrix organization. Besides, we found significant differences in the expression of SNAI2 and Prrx1, which may be one of the reasons for the changes of EMT (Supplementary File S4). Extracellular matrix becomes highly dysregulated in cancer, playing both protumorigenic and antitumorigenic roles. Ovarian cancer cell injection may induce changes of extracellular stromal environment in pancreatic cancer and then lead to the changes of cell adhesion, differentiation, migration, proliferation and invasion in pancreatic cancer ^33^. It is worth noting that many cancers cause death because of cancer recurrence and metastasis. Ovarian cancer cell treatment for pancreatic cancer could cause a reduction in the invasion and migration ability of pancreatic cancer, such an effect may be a new breakthrough in the clinical treatment of cancer.

Immunotherapy achieved remarkable efficacy in many malignancies ^34–38^, but has not yet translated to pancreatic cancer. The failure of immunotherapy is mainly due to the stromal microenvironment within the tumor that affects the activity of immune checkpoint blockers ^34,39^ and the infiltration of CAR-T cells into the tumor ^40–42^. The CCC therapeutics does not alter the immune pathways, so it may do not belong to immunotherapy. This study may open up a new type of therapeutics for cancer treatment, not only including pancreatic cancer but also perhaps including more cancers in the future, such as glioblastoma, the most lethal cancer in central nervous system.

In addition, it is interesting to discover some potential drugs for the treatment of pancreatic cancer by analyzing the RNA-seq data from the CCC therapeutics based on the drug related categories such as ‘Drug Perturbations from GEO’ and “DSigDB”. As a result, 120 connections between drugs and these DEGs were identified as significance (Supplementary File S6). It should be noted that these ‘connection’ identified here is association, therefore, whether it alleviate or enhance pancreatic cancer still need expert decision and further experiments. After manual curation, we further found 113 of the 120 connections have clear evidence that corresponding drugs can treat cancer (Supplementary File S6), suggesting that they may be also effective for treating pancreatic cancer.

Obviously, a number of scientific and technique questions need to be further answered. Firstly, the feasibility of intratumoral injection in clinical practice remains to be explored. For clinics, surgery with translation of the therapeutic cancer cells into the tumor would be an option. Secondly, in this study, only two doses were selected for the animal experiments. The dose of injected cells needs to be further explored in more details to achieve better efficacy. In addition, the exact mechanisms of the CCC therapeutics still need to be explored. Moreover, only tumor weight and tumor volume were investigated in the present study, survival time is also important to be explored in the future. Next, more germline of cell lines should be tested in the validation experiments. The ovarian cancer cell lines include CAOV3, PA1 and SKOV3 and the pancreatic cancer cell lines include Capan-1/2, SW1990, CFPAC-1, HPAC, HPAF-2, Hs-766T, PANC-1, and so on. At present study, we only tested SKOV3 as the therapeutic cancer cell and SW1990 as the xenograft cell of the pancreatic tumor. It is thus quite important to investigate the efficacy of the CCC therapeutics using more types of cancer cells. Finally, it is also very important to test whether the CCC therapeutics is a general strategy for the treatment of other cancers.

## Credit authorship contribution statement

Q.C. proposed the original idea. Q.C., X.L. and C.C. designed the work flow of the experiments. X.L. performed the experiments. C.C. analyzed the data. X.L. wrote the draft manuscript. Q.C., X.L., and C.C. wrote and finalized the manuscript.

## Acknowledgments

We thank Prof. Lixiang Xue for providing the SKOV3 cell line and thank Co., Ltd of Beijing JeaMoon Technology for the DrugMine tool.

## Reference

1. Integrated Genomic Characterization of Pancreatic Ductal Adenocarcinoma. Cancer Cell. 2017;32: 185–203.e113.

2. Bray F, Ferlay J, Soerjomataram I, Siegel RL, Torre LA, Jemal A. Global cancer statistics 2018: GLOBOCAN estimates of incidence and mortality worldwide for 36 cancers in 185 countries. CA Cancer J Clin. 2018;68: 394–424.

3. Rahib L, Smith BD, Aizenberg R, Rosenzweig AB, Fleshman JM, Matrisian LM. Projecting cancer incidence and deaths to 2030: the unexpected burden of thyroid, liver, and pancreas cancers in the United States. Cancer Res. 2014;74: 2913–2921.

4. Strobel O, Neoptolemos J, Jäger D, Büchler MW. Optimizing the outcomes of pancreatic cancer surgery. Nat Rev Clin Oncol. 2019;16: 11–26.

5. Auclin E, Marthey L, Abdallah R, et al. Role of FOLFIRINOX and chemoradiotherapy in locally advanced and borderline resectable pancreatic adenocarcinoma: update of the AGEO cohort. Br J Cancer. 2021;124: 1941–1948.

6. Ahmad SA, Duong M, Sohal DPS, et al. Surgical Outcome Results From SWOG S1505: A Randomized Clinical Trial of mFOLFIRINOX Versus Gemcitabine/Nab-paclitaxel for Perioperative Treatment of Resectable Pancreatic Ductal Adenocarcinoma. Ann Surg. 2020;272: 481–486.

7. Conroy T, Hammel P, Hebbar M, et al. FOLFIRINOX or Gemcitabine as Adjuvant Therapy for Pancreatic Cancer. N Engl J Med. 2018;379: 2395–2406.

8. Conroy T, Desseigne F, Ychou M, et al. FOLFIRINOX versus gemcitabine for metastatic pancreatic cancer. N Engl J Med. 2011;364: 1817–1825.

9. Von Hoff DD, Ervin T, Arena FP, et al. Increased survival in pancreatic cancer with nab-paclitaxel plus gemcitabine. N Engl J Med. 2013;369: 1691–1703.

10. Nevala-Plagemann C, Hidalgo M, Garrido-Laguna I. From state-of-the-art treatments to novel therapies for advanced-stage pancreatic cancer. Nat Rev Clin Oncol. 2020;17: 108–123.

11. Larkin J, Chiarion-Sileni V, Gonzalez R, et al. Five-Year Survival with Combined Nivolumab and Ipilimumab in Advanced Melanoma. N Engl J Med. 2019;381: 1535–1546.

12. Dummer R, Lebbé C, Atkinson V, et al. Combined PD-1, BRAF and MEK inhibition in advanced BRAF-mutant melanoma: safety run-in and biomarker cohorts of COMBI-i. Nat Med. 2020;26: 1557–1563.

13. Rittmeyer A, Barlesi F, Waterkamp D, et al. Atezolizumab versus docetaxel in patients with previously treated non-small-cell lung cancer (OAK): a phase 3, open-label, multicentre randomised controlled trial. Lancet. 2017;389: 255–265.

14. Ansell SM, Lesokhin AM, Borrello I, et al. PD-1 blockade with nivolumab in relapsed or refractory Hodgkin’s lymphoma. N Engl J Med. 2015;372: 311–319.

15. Reck M, Rodríguez-Abreu D, Robinson AG, et al. Pembrolizumab versus Chemotherapy for PD-L1-Positive Non-Small-Cell Lung Cancer. N Engl J Med. 2016;375: 1823–1833.

16. Ascierto PA, Avallone A, Bhardwaj N, et al. Perspectives in Immunotherapy: meeting report from the Immunotherapy Bridge, December 1st-2nd, 2021. J Transl Med. 2022;20: 257.

17. Wu AA, Bever KM, Ho WJ, et al. A Phase II Study of Allogeneic GM-CSF-Transfected Pancreatic Tumor Vaccine (GVAX) with Ipilimumab as Maintenance Treatment for Metastatic Pancreatic Cancer. Clin Cancer Res. 2020;26: 5129–5139.

18. Brown CE, Alizadeh D, Starr R, et al. Regression of Glioblastoma after Chimeric Antigen Receptor T-Cell Therapy. N Engl J Med. 2016;375: 2561–2569.

19. O’Rourke DM, Nasrallah MP, Desai A, et al. A single dose of peripherally infused EGFRvIII-directed CAR T cells mediates antigen loss and induces adaptive resistance in patients with recurrent glioblastoma. Sci Transl Med. 2017;9.

20. Tchou J, Zhao Y, Levine BL, et al. Safety and Efficacy of Intratumoral Injections of Chimeric Antigen Receptor (CAR) T Cells in Metastatic Breast Cancer. Cancer Immunol Res. 2017;5: 1152–1161.

21. Siegel RL, Miller KD, Fuchs HE, Jemal A. Cancer Statistics, 2021. CA Cancer J Clin. 2021;71: 7–33.

22. Lamb J, Crawford ED, Peck D, et al. The Connectivity Map: using gene-expression signatures to connect small molecules, genes, and disease. Science. 2006;313: 1929–1935.

23. Haider S, Wang J, Nagano A, et al. A multi-gene signature predicts outcome in patients with pancreatic ductal adenocarcinoma. Genome Med. 2014;6: 105.

24. Sherman BT, Hao M, Qiu J, et al. DAVID: a web server for functional enrichment analysis and functional annotation of gene lists (2021 update). Nucleic Acids Res. 2022;50: W216–221.

25. Fan R, Cui Q. Toward comprehensive functional analysis of gene lists weighted by gene essentiality scores. Bioinformatics. 2021.

26. Cobo I, Martinelli P, Flández M, et al. Transcriptional regulation by NR5A2 links differentiation and inflammation in the pancreas. Nature. 2018;554: 533–537.

27. Hu D, Ansari D, Zhou Q, Sasor A, Said Hilmersson K, Andersson R. Stromal fibronectin expression in patients with resected pancreatic ductal adenocarcinoma. World J Surg Oncol. 2019;17: 29.

28. Hollstein M, Sidransky D, Vogelstein B, Harris CC. p53 mutations in human cancers. Science. 1991;253: 49–53.

29. Abegglen LM, Caulin AF, Chan A, et al. Potential Mechanisms for Cancer Resistance in Elephants and Comparative Cellular Response to DNA Damage in Humans. Jama. 2015;314: 1850–1860.

30. Nieto MA, Cano A. The epithelial-mesenchymal transition under control: global programs to regulate epithelial plasticity. Semin Cancer Biol. 2012;22: 361–368.

31. Valastyan S, Weinberg RA. Tumor metastasis: molecular insights and evolving paradigms. Cell. 2011;147: 275–292.

32. Bakiri L, Macho-Maschler S, Custic I, et al. Fra-1/AP-1 induces EMT in mammary epithelial cells by modulating Zeb1/2 and TGFβ expression. Cell Death Differ. 2015;22: 336–350.

33. Cox TR. The matrix in cancer. Nat Rev Cancer. 2021;21: 217–238.

34. Brahmer JR, Tykodi SS, Chow LQ, et al. Safety and activity of anti-PD-L1 antibody in patients with advanced cancer. N Engl J Med. 2012;366: 2455–2465.

35. Sharma P, Callahan MK, Bono P, et al. Nivolumab monotherapy in recurrent metastatic urothelial carcinoma (CheckMate 032): a multicentre, open-label, two-stage, multi-arm, phase 1/2 trial. Lancet Oncol. 2016;17: 1590–1598.

36. Motzer RJ, Escudier B, McDermott DF, et al. Nivolumab versus Everolimus in Advanced Renal-Cell Carcinoma. N Engl J Med. 2015;373: 1803–1813.

37. Robert C, Long GV, Brady B, et al. Nivolumab in previously untreated melanoma without BRAF mutation. N Engl J Med. 2015;372: 320–330.

38. Borghaei H, Paz-Ares L, Horn L, et al. Nivolumab versus Docetaxel in Advanced Nonsquamous Non-Small-Cell Lung Cancer. N Engl J Med. 2015;373: 1627–1639.

39. Le DT, Picozzi VJ, Ko AH, et al. Results from a Phase IIb, Randomized, Multicenter Study of GVAX Pancreas and CRS-207 Compared with Chemotherapy in Adults with Previously Treated Metastatic Pancreatic Adenocarcinoma (ECLIPSE Study). Clin Cancer Res. 2019;25: 5493–5502.

40. Vonderheide RH, Bayne LJ. Inflammatory networks and immune surveillance of pancreatic carcinoma. Curr Opin Immunol. 2013;25: 200–205.

41. Johnson BA, 3rd, Yarchoan M, Lee V, Laheru DA, Jaffee EM. Strategies for Increasing Pancreatic Tumor Immunogenicity. Clin Cancer Res. 2017;23: 1656–1669.

42. Stromnes IM, Hulbert A, Pierce RH, Greenberg PD, Hingorani SR. T-cell Localization, Activation, and Clonal Expansion in Human Pancreatic Ductal Adenocarcinoma. Cancer Immunol Res. 2017;5: 978–991.

